# Effects of C-peptide replacement therapy on bone microarchitecture parameters in streptozotocin-diabetic rats

**DOI:** 10.1101/2020.01.27.921288

**Authors:** Samantha Maurotti, Cristina Russo, Vincenzo Musolino, Saverio Nucera, Micaela Gliozzi, Miriam Scicchitano, Francesca Bosco, Valeria Maria Morittu, Monica Ragusa, Elisa Mazza, Carmine Gazzaruso, Domenico Britti, Maria Teresa Valenti, Michela Deiana, Stefano Romeo, Sandro Giannini, Luca Dalle Carbonare, Vincenzo Mollace, Arturo Pujia, Tiziana Montalcini

**Affiliations:** Department of Medical and Surgical Science, Magna Græcia University of Catanzaro, Italy; Department of Clinical and Experimental Medicine, Magna Græcia University of Catanzaro, Catanzaro, Italy; IRC-FSH Interregional Center for Food Safety and Health, Department of HealthSciences, Magna Græcia University of Catanzaro, Italy; Department of Health Sciences, Magna Græcia University of Catanzaro, Italy; Diabetes and endocrine and metabolic diseases Unit and the Centre for Applied Clinical Research (Ce.R.C.A.) Clinical Institute “Beato Matteo” (Hospital Group San Donato), Vigevano, Italy; Department of Medicine, Specialized Regional Center for Biomolecular and Histomorphometric Research on Degenerative and Skelatal Diseases, Verona, Italy; Department of Neurosciences, Biomedicine and Movement Sciences, University of Verona, Italy; Department of Molecular and Clinical Medicine, Sahlgrenska Center for Cardiovascolar and Metabolic Research, University of Gothenburg, Göteborg, Sweden; Department of Medicine, Clinica Medica 1, University of Padova and Regional Center for Osteoporosis, Padova, Italy

**Keywords:** c-peptide, bone microarchitectural parameters, Type 1 diabetes mellitus, osteoporosis, diabetic rat model

## Abstract

Pathological pathways involved in the development of diabetic complications are still unclear. Several studies suggested a pathogenic role of C-peptide deficiency in vascular and neuropathic complications. However, to date, the role of C-peptide on diabetes-related bone loss has not been investigated. The objective of this study was to test the effects of a 6-week regimen of rat C-peptide infusion on bone by combining micro-CT imaging with histological testing.

Twenty-three, four-month-old male Wistar rats were randomly divided into three groups: Normal control group; Sham diabetic control group; Diabetic plus C-peptide group. Diabetes was induced by injection of streptozotocin. Rat C-peptide was delivered via subcutaneously osmopumps. We assessed several trabecular microarchitectural parameters and cellular and matrix proteins in bone tissue sections.

At the end of the study, both the normal control and diabetic plus C-peptide groups had a higher tibia weight than the diabetic control group (*p* = 0.05 and *p* = 0.02, respectively). C-peptide levels significantly and positively correlated with the trabecular thickness (r= 0.45, p=0.08) and negatively with structure model index (r= −0.40, p=0.09). Furthermore, it positively correlated with one of the histological assessments (i.e. PLIN1 score; p=0.02). Micro-CT evaluation showed significant inter-group differences in trabecular thickness (p=0.02), trabecular space (*p* = 0.05) and cross-sectional thickness (p=0.05). Diabetic plus C-peptide group showed a higher trabecular thickness (p<0.001), cross-sectional thickness (p=0.03) and a lower trabecular space (p=0.05) than the normal control group. Both the normal control and diabetic plus C-peptide groups had more Runx-2 and PLIN1 positive cells in comparison to the diabetic control group (p = 0.045 and p = 0.034).

For the first time we demonstrated that the diabetic rats receiving rat C-peptide had higher quality of trabecular bone, than diabetic rats not receiving it. C-peptide could have a role in the prevention of diabetes-related bone loss.

## 1. Introduction

While new developments have been shown to improve acute life-threatening complications in patients with type 1 diabetes mellitus (T1DM) [1], long-term complications are still an ongoing burden for these patients [2]. Unfortunately, T1DM complications develop even in individuals with good glycemic control [3]. The risk of complications may, thus, be mainly linked to the duration of the disease [4].

However, this is not the case for bone loss in T1DM. A decreased bone mass has been reported for prepubertal and pubertal patients with T1DM [5]. Bone loss has been found in over 50% of patients with juvenile-onset diabetes [6,7], and it is worrying that fractures are associated with a disproportionally high mortality rate in children [8,9]. In contrast, the prevalence of cardiovascular disease among T1DM patients increases with age, from only 6% in patients aged 15-29 years to 25% in patients aged 45-59 years [10]. It has thus been suggested that adverse effects on bone health may occur earlier after a diabetes diagnosis than subclinical cardiovascular adverse effects [11–13]. Of note, bone loss develops even in individuals under insulin therapy [14]. Thus, neither the disease duration nor the glycaemic control explain the enigma of diabetic complications.

Insights into the possible mechanisms leading to the development of diabetic complications were gained by the finding that C-peptide influences several clinical outcomes in diabetic patients.

C-peptide is the 31-amino acid segment of the proinsulin hormone, which is secreted in equimolar amounts with insulin from the pancreatic β-cells, remaining intact after cleavage from insulin [15, 16]. C-peptide replacement therapy has been studied in several diabetic animal models as well as human studies which show consistent beneficial effects against the development of microvascular and neuropathic diabetes complications [17–24].

One study demonstrated that C-peptide influences in vitro the biological behavior of human osteoblast-like cells [25]. However, to date, the effects of C-peptide replacement therapy on bone in T1DM has not been investigated either in *in vivo* or clinical studies.

Recently, two cross-sectional studies suggested a pathogenic role of C-peptide deficiency in T1DM-related osteoporosis [26,27].

The current work thus represents the first study investigating the effects of C-peptide replacement therapy on bone in diabetic rats. The most commonly used T1DM animal model is the streptozotocin (STZ) rodent model [17]. Rapid bone loss also occurs in STZ-induced diabetic rats [28,29], and, over time, this model has been widely used for studying osteoporosis.

Our study assessed the effects of a 6-week continuous administration of rat C-peptide on bone in STZ-induced diabetic rats.

These results may serve as a basis for future human studies on the effects of C-peptide therapy in patients affected by osteoporosis.

## 2. Materials and Methods

### 2.1 Ethical approval

The experiments were carried out following the European guidelines (2010/63/EU) regarding procedures with animals used, in accordance with the approval of the Ethics Committee for Experimental Animals Welfare of the University Magna Grecia, Catanzaro (auth. 10/01/2018) and Italian Ministry of Health (auth. N° 353/2018-protocol ADEAB.16, auth. 9/05/2018).

### 2.2 Animals

Twenty-three, four-month-old male Wistar rats (Charles River Laboratories) (400–500 g) were used. The rats were fed with a commercial standard diet containing 1 % calcium, 0.7 % phosphorus (of which 0.4 % non-phytate phosphorus), Ca/P 3:1 and 150 IU of vitamin D3 per 100 g. All rats were individually housed in polycarbonate cages and were kept under the same conditions. The environmental conditions were maintained constant with 12 h/12 h light/day cycle, 24 ± 1 °C temperature and 55 ± 10% humidity. The animals had ad libitum access to pellets of standard rodent diet, as well as bottles of tap water.

### 2.3 Experimental Design

The rats were randomly divided into the following 3 groups:

-Normal control group (no treatment; sacrificed at the end of the experiment)
-Diabetic control group (no treatment; sacrificed at the end of the experiment)
-Diabetic plus C-peptide group (C-peptide-treated; sacrificed at the end of the experiment)

Animal groups are abbreviated as follows:

-Normal control group= CTR
-Diabetic control group=D-CTR
-Diabetic plus C-peptide group=C-PEP

After a 2-week acclimation period, diabetes was induced by a single intraperitoneal injection of STZ (60 mg/kg body weight) dissolved in 0.05 M citric acid (pH 5.1). One week after STZ injection, serum was collected by tail bleeding, and the rats with serum glucose concentrations greater than 300 mg/dl (16.6 mmol/L) throughout a four weeks period were regarded as diabetic. To prevent ketosis, a moderate dose of rapid and intermediate insulin was administered (Humalog and Humulin I, Eli Lilly Italia S.p.A.) when blood sugar levels were higher than 400 mg/dl in order to maintain a glucose level below 350 mg/dl. A total of 3 rats died after diabetes induction.

In C-PEP group, the rat C-peptide (>95% purity, RP-HPLC) was delivered via subcutaneously implanted osmopumps (Alzet 2006, Alza, Palo Alto, CA) delivering 72 nmol • kg^−1^ • 24 h^−1^ [18] for 6 weeks [28], while the D-CTR group was sham-operated. Rat C-peptide was provided by Dr John Wahren, Karolinska Institutet, Stockholm, Sweden.

During the course of this study, animals were weighed weekly and nonfasting blood glucose levels were recorded every day. The animals were euthanized after 6 weeks (bone depletion period) [28] by isoflurane inhalation.

The study design is outlined in Figure 1.

**Figure 1:**
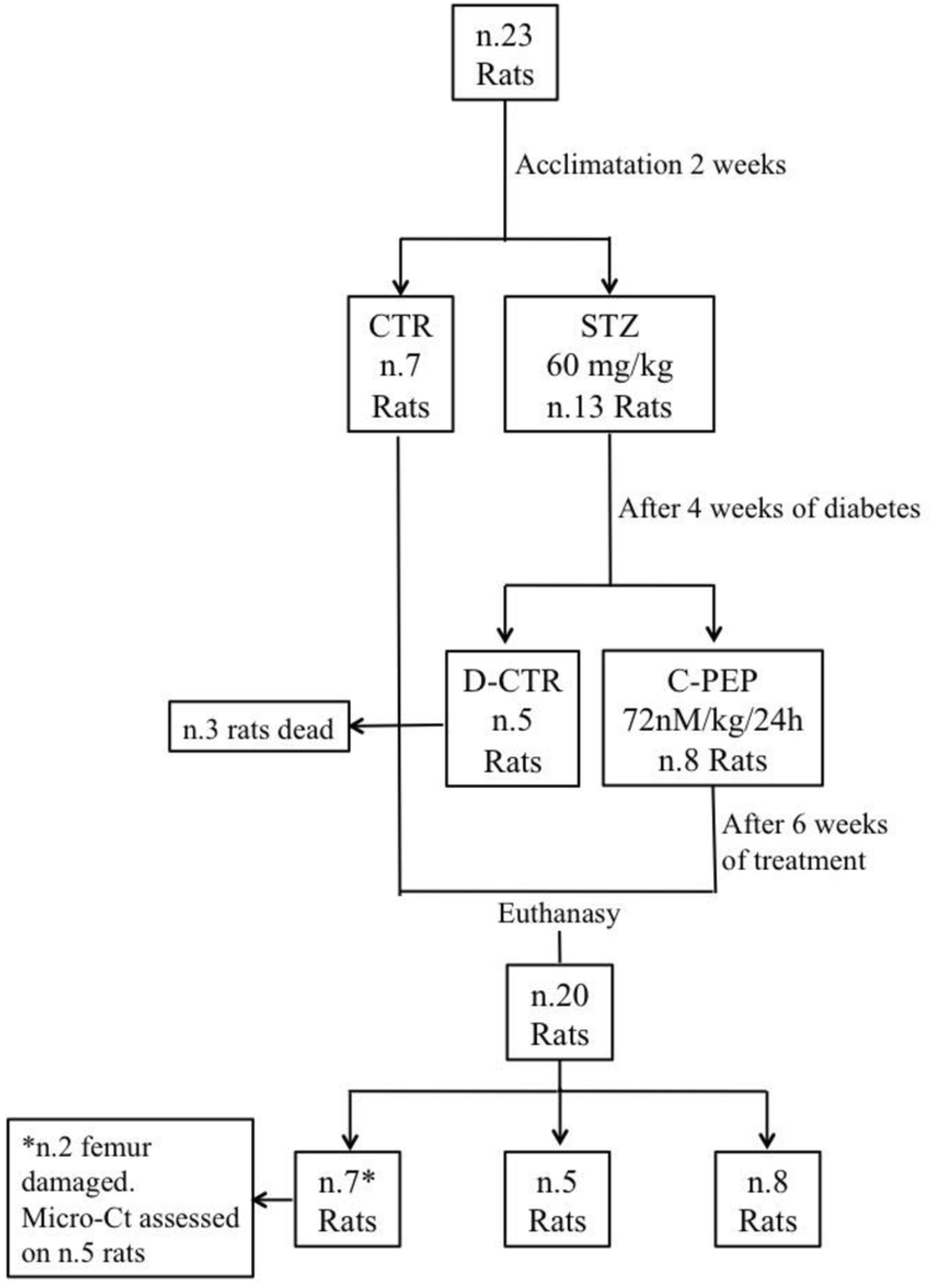
Study design

### 2.4 Tissue collection

Femurs and tibiae were obtained postmortem, dissected and cleaned of adherent soft tissues.

The right femurs were immediately scanned using micro-computed tomography (micro-CT) as described below. The left femurs were fixed by immersion in formalin and stored for histologic assessments. The left tibiae from each animal were stored in plastic vials at −70°C for ashing to assess calcium content.

### 2.5 Parameters of general health and bone metabolism

Venous blood was collected at baseline (the first day of C-peptide delivery) and at end of the study and serum was stored frozen at −20°C until assayed.

Specifically, serum glucose, calcium and phosphate concentration were analysed by Roche Module COBAS 8000 (Roche Diagnostics, Indianapolis, IN).

Serum C-peptide was assessed with an enzymatic colorimetric test by ELISA kit (Crystal chem, Inc. USA). Both the CTX and RANKL were measured by ELISA kits (Cusabio Biotech Co, USA) but RANKL was measured only at the end of the study.

The haematological parameters were analysed by using an automated hematological analyser (ADVIA Siemens Healthcare Diagnostic Inc.; Germany). All measurements were performed in duplicate.

### 2.6 Micro-computed tomography

We scanned the samples using Skyscan 1176 micro-CT system (Bruker microCT, Skyscan, Belgium). In order to avoid drying of the samples, we wrapped the femur of each animal with a thin layer paper soaked in saline solution and then analysed along the longitudinal axis of the specimen [30].

We analysed the distal femur’s portion [31]. under the following scanning conditions and parameters: 65 kV, 380 μA, 25 W, 950 ms exposure time, 1 mm Aluminum filter, 1×1 camera binning and 9 μm isotropic resolution, rotation step of 0.3°, 12.47 um camera pixel size.

The reconstruction of the projection images was performed in NRecon software (Bruker microCT, Belgium) on InstaRecon platform (InstaRecon, USA) with appropriate thermal drift correction, misalignment compensation, Gaussian smoothing of 1, and appropriate ring artifact and beam hardening corrections. Ct.An program version 1.15.4 (Skyscan, Belgium) was used for quantitative analysis of the data. The volume of interest (VOI) was identified as follow: in each femur the reference slice corresponding to the upper end of the distal femur’s growth plate cartilage was identified, according to the manufacturer’s recommendation, and trabecular VOI comprised the slices from number 100 to 500 proximal to the reference slice. Hand-drawn contours were used to isolate the metaphyseal region of interest and trabecular compartments based on 400 consecutive slices [32].Threshold was set automatically through Ct.An software (Otsu automatic threshold); subsequently, the suggested values were checked manually by simultaneous overview of raw and thresholded images, which allowed us to fine-tune the threshold level to best represent the mineralized bone tissue. All the procedures were standardised and the same threshold was applied for all specimens to allow inter-individual comparisons of microarchitectural parameters.

[31]. Trabecular parameters were as follows: Trabecular volumetric bone mineral density [vBMD; mg hydroxyapatite (HA)/cm^3^], bone volume fraction (BV/TV; %), trabecular thickness (Tb.Th; mm), trabecular number (Tb.N; 1/mm), trabecular separation (Tb.Sp; mm), structure model index (SMI), connectivity density (Conn.D; 1/mm^3^) and cross-sectional thickness (Cs.Th; mm).

The parameters of bone geometry were also assessed: bone perimeter (B.Pm, mm) that represents total perimeter of the diaphyseal bone cross-section, and tissue perimeter (T.pm; mm).

### 2.7 Histomorphometry

The study of microarchitecture was based on the measure of width, number, and separation of trabeculae as well as on their spatial organization.

The samples were fixed in 70% ethanol and embedded undecalcified in methyl-methacrylate resin. Bone sections were cut by using a microtome (Pfm Rotary 3000 Bio-Optica, Germany) equipped with a carbide-tungsten blade, stained with Goldner’s stain and mounted on microscope slides for histomorphometric measurements. Measurements were performed by means of an image analysis system consisting of an epifluorescent microscope (Leica DM2500) connected to a digital camera (Leica DFC420 C) and a computer equipped with a specific software for histomorphometric analyses (Bone 3.5, Explora Nova, France). We evaluated parameters of structure such as bone volume/tissue volume (BV/TV; %), trabecular thickness (Tb.Th; mm), trabecular number (Tb.N; N/mm), trabecular separation (Tb.Sp; mm); and of microarchitecture such as marrow star volume (MSV, N/mm^3^), Trabecular Bone Pattern Factor (TBPf) and fractal dimension (FD) as previously described [33,34]. Histomorphometric parameters were reported in accordance with the ASBMR Committee nomenclature [35].

### 2.8 Immunohistochemistry

In order to identifies the proteins in tissue sections (cellular and matrix components of bone), samples from femur mid-diaphysis embedded in paraffin underwent immunohistochemistry (IHC) analyses by incubating the samples with antibodies specific to the protein of interest, and then visualizing the bound antibody using a chromogen [36].

Osteoblast differentation in decalcified bone sections of the rats was investigated in transverse sections (5 μm) from femur by incubating the samples with mouse anti-Runx-2 (an osteoblast marker) antibody (1:1000 dilution in PBS containing 0.5% BSA, Cell Signaling Technology) at 4 °C. Sections were washed with PBS and signal was detected using the Ultravision ONE Detection System HRP polimer & DAB Plus Chromogen (Thermo Scientific) with Runx-2 positive cells were counted in five randomly selected fields from six bone sections of each group and quantified with Image-Pro plus 6.1 software.

Since there could be an associations between proteins that regulate fat metabolism and skeletal mass, immunohistochemical staining for perilipin 1 (PLIN1, which coats the lipid droplets in adipocytes) [37] was performed incubating the samples with mouse anti-PLIN1 antibody (1:1000 diluition in PBS containing 0.5% BSA, Cell Signaling Technology).

An immunohistochemical analysis was also conducted to investigate on the proteins eventually involved in tensile strength and bone healing by monoclonal antibodies against collagen I alpha 1 (COLIA1, 1:500, Santa Cruz biotechnology) and bone sialoprotein (BSP, 1:1000 diluition in PBS containing 0.5% BSA, Cell Signaling Technology).

We count the percentage of positive immunolabeled cells over the total cells in each selected area.

### 2.9 Anthropometric and Compositional analysis of the tibia (calcium content)

Left and right tibiae were dissected and weighed to 0.1 g accuracy on an electronic balance. A digital caliper accurate to 0.01 mm was used to measure the tibiae length in the sagittal plane. These measurements were carried out three times and the average was calculated. We then calculated the mean between left and right tibiae.

The left tibiae were dried in a vacuum oven at 110 °C for 6 hours and the dry mass was recorded. Then tibiae were ashed at 800 °C for 4 h, weighed again and dissolved in 1 mL 6N HCl. Calcium content (mg/dl) was obtaining by colorimetric determination with Quantichrom calcium Assay kit [38] (BioAssay Systems, Hayward, CA).

### 2.10 Statistical Analysis

The results are expressed as mean ± S.D.

Sample size was calculated with G*Power software, version 3.1.9.2. A minimum sample size estimate of 5 animal per group was required to obtain 80 % power to detect a maximum difference of 2% in the Tb.Th mean between control and C-peptide group (not inferior), using a two-sided 0.05 significance level. We calculated an effect size equal to 0.0024 (0.122*0.02), with an effect size / standard deviation (E/SD) ratio equal to 2.44.

Continuous data normalcy was analysed through the Shapiro-Wilk test and Box plot method. The outlier values that were considered due to measurement error or capture were excluded from the analysis. In the case of normal data, ANOVA test was used to compare the means between groups (specifically, anthropometric and laboratory parameters). All the significant differences were adjusted for the baseline body weight by a General Linear Model (GLM) with Bonferroni correction followed by LSD test as *post hoc* analysis. In the case of non-normal data (microarchitectural parameters and RANKL), the nonparametric Kruskal-Wallis followed by Mann-Whitney tests were used to evaluate the differences between groups. All the significant differences were adjusted for the baseline body weight by an ordinal regression analysis using only categorical dependent variables. Changes from baseline to the end of the study (within group variation in body weight, glucose, electrolytes and C-peptide) were assessed using a paired Student’s *t*-test (two tailed).

We categorized the immunohistochemistry scores (of PLIN1, RUNX, BSPI and COL1A) according to three groups of values (group I, score <3; group II, score equal to 3; group III, score >3) and a Chi-squared test was used for comparisons of these categorical variables.

Pearson and Spearman’s correlation was performed for normal and non-normal data, respectively, to assess the statistical dependence between variables. A probability value of less than 0.05 was considered statistically significant. All statistical analyses were performed using SPSS for Windows, version 22.0 (IBM Corporation, New York, NY, United States).

## 3. Results

### 3.1 General Health

At baseline, the CTR group body weight was significantly lower than D-CTR and C-PEP body weights (*p* < 0.005 and *p* < 0.030, respectively, Table 1). The CTR rats gained weight during the experimental period of 42 days, whereas the D-CTR rats had 11% and the C-PEP group had 7% decreases in the final body weight, compared to the initial weight (Table 1).

**Table 1:** Baseline and final clinical characteristics of rats according to intervention group

| Variables | CTR<sup>a</sup><br>(n=5) | D-CTR<sup>b</sup><br>(n=5) | C-PEP<sup>c</sup><br>(n=8) | *p*<sup>*</sup> | *Post-hoc* | CTR<sup>a</sup><br>(n=5) | D-CTR<sup>b</sup><br>(n=5) | C-PEP<sup>c</sup><br>(n=8) | *p*<sup>*</sup> | *Post-hoc* |
| --- | --- | --- | --- | --- | --- | --- | --- | --- | --- | --- |
|  | *Basal* |  |  |  |  | *Final* |  |  |  |  |
| Body weight (g) | 488±48 | 554±43 | 569±54 | 0.016 | c vs a, p=0.030<br>b vs a, p=0.005 | 540±30 | 493±38 | 529±33 | 0.097 | b vs a, p=0.04 |
| Glucose (mg/dL) | 120±21 | 111±17 | 105±14 | 0.31 | / | 102±23 | 273±4 | 115±53 | 0.012 | b vs a, p=0.08<br>c vs b, p=0.08 |
| Calcium (mg/dL) | 9.9 ±0.2 | 10.5±0.1 | 10.3±0.1 | 0.22 | / | 10.6±0.1 | 10.3±0.3 | 10.8±0.3 | 0.042 | b vs c, p=0.04 |
| Phosphorus (mg/dL) | 6.82±0.2 | 7.19±0.2 | 7.5±0.2 | 0.17 | / | 6.0±0.6 | 6.4±0.4 | 7.3±0.4 | 0.01 | b vs c, p=0.05<br>a vs c, p=0.02 |
| Creatinine (mg/dL) | 0.33±0.03 | 0.37±0.04 | 0.45±0.01 | 0.33 | / | 0.40±0.02 | 0.46±0.02 | 0.39±0.02 | 0.29 | / |
| C-peptide (mg/dL) | 1.54±1 | 1.95±0.1 | 1.52±0.4 | 0.51 | / | 1.58±0.1 | 0.44±0.1 | 0.98±0.1 | 0.002 | a vs c, p=0.09<br>b vs a, p=0.05<br>b vs c, p=0.02 |
| CTX-I (pg/mL) | 886±100 | 826±85 | 820±67 | 0.33 | / | 802±103 | 914±109 | 801±83 | 0.11 | / |
| Rapid Insulin (U/day) | / | / | / | / | / | / | 1.4±1.7 | 1.4±2 | 0.90 | / |
| Intermediate Insulin (U/day) | / | / | / | / | / | / | 2.2±3 | 2±3 | 0.28 | / |

Note. Data are mean ± SD. Each significative difference is adjusted for the baseline body weight (except glucose); CTX-I C-telopeptide of type I collagen. <sup>\*</sup>Unpaired t-test.

STZ injection resulted in a significant reduction of serum C-peptide in both the sham D-CTR and the C-PEP group relative to the CTR group, at the end of the treatment phase (Figure 2). The D-CTR rats had 79% and the C-PEP group had 41% decreases in the final serum C-peptide compared to the initial C-peptide level (Fig. 2). At the end of the study, serum C-peptide was higher in both the CTR (*p* = 0.05) and C-PEP groups (*p* = 0.02) than in the D-CTR group, whereas C-peptide levels did not differ between CTR and C-PEP groups (*p* = 0.09) (Table 1; Figure 2D).

**Figure 2:**
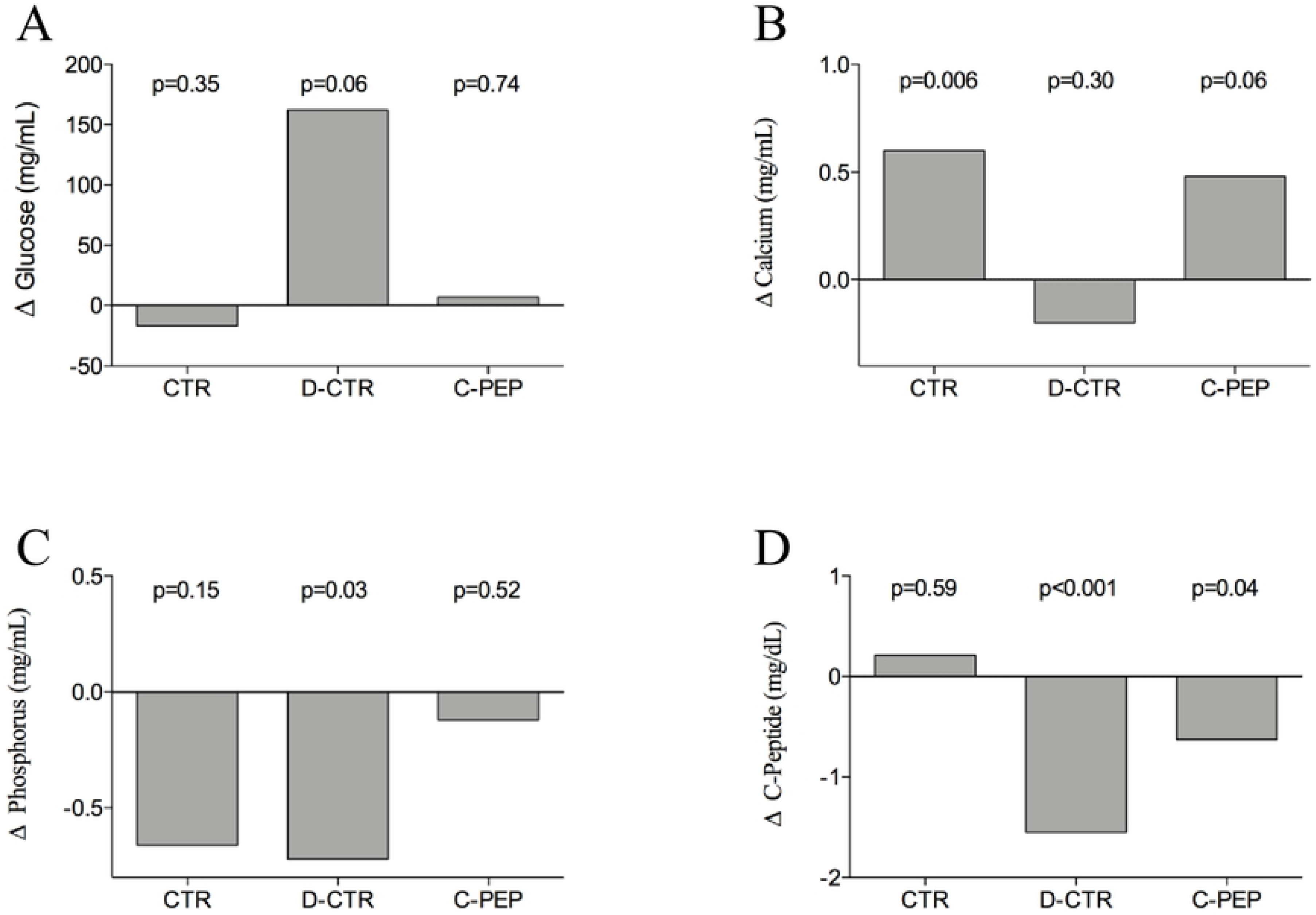
Change in laboratory parameters at paired t-test according to intervention groups of serum (A) glucose (B) calcium (C) phosphorus and (D) C-Peptide.

Moreover, at the end of the study, the C-PEP group had a higher serum phosphorus level than the CTR (p=0.02) and the D-CTR (p=0.05) groups as well as a higher calcium levels than the D-CTR (p=0.04). Considering the results of the paired t-test, the phosphorus variations were: −0.66, −0.72 and −0.12 in the CTR, D-CTR and C-PEP groups, respectively (p=0.15, p=0.03; p =0.52, respectively; Table 1, Figure 2C).

C-peptide infusion prevented the reduction of red cells and haemoglobin seen in the D-CTR group (for red cells, D-CTR vs C-PEP, p < 0.004; for haemoglobin, D-CTR vs C-PEP, p < 0.005; Table 2).

**Table 2:** Clinical characteristics of rats assessed at the end of the study (only once), according to intervention group

| Variables | CTR<sup>a</sup><br>(n=5) | D-CTR<sup>b</sup><br>(n=5) | C-PEP<sup>c</sup><br>(n=8) | *p*-value | *p*-value<br>*post hoc* |
| --- | --- | --- | --- | --- | --- |
| Femur weight (g)* | 1.46±0.07 | 1.54 ±0.17 | 1.55±0.09 | 0.41 | / |
| Femur length (cm)* | 4.5±0.2 | 4.8±0.1 | 4.5±0.1 | 0.49 | / |
| Tibia weight (g)* | 1.2±0.08 | 1.06±0.05 | 1.2±0.05 | 0.09 | c vs b, p=0.02<br>a vs b, p=0.05 |
| Tibia length (g)* | 5±0.2 | 5.3±0.1 | 5.2±0.1 | 0.69 | / |
| Tibia calcium content (g/dL) | 5.8±6 | 3.18±5 | 5.7±6 | 0.75 | / |
| Ash weight (g) | 0.38±0.03 | 0.39±0.05 | 0.45±0.03 | 0.022 | a vs c, p=0.016<br>b vs c, p=0.02 |
| RBC (10<sup>6</sup>/μL) | 9.1±0.2 | 8.6±0.7 | 9.8 ±0.7 | 0.01 | b vs c, p= 0.004 |
| Hemoglobin (g/dL) | 15.5±0.3 | 13.8±1 | 16.1±1 | 0.019 | b vs c, p= 0.006 |
| WBC (10<sup>3</sup>/μL) | 6.5±3 | 11.4±3 | 11.8±0.3 | 0.131 | / |

Note. Data are mean ± SD. \*mean value between left and right; each significative difference is adjusted for the baseline body weight;; RBC Red Blood Cell; WBC White Blood Cell

### 3.2 Anthropometric and Compositional analysis of the tibia

At the end of the study, both the CTR and C-PEP groups exhibited a higher mean tibia weight than the D-CTR group (CTR vs D-CTR, *p* = 0.05 and C-PEP vs D-CTR, *p* = 0.02, Table 2). Calcium content did not differ between groups while the ash weight was higher in C-PEP than in the other groups (p=0.022; Table 2).

### 3.3 Correlation between serum C-peptide, RANKL and microarchitecture parameters

S1 Table shows a correlation between several radiological (micro-CT) parameters of microarchitecture (such as Tb.Th and Cs.Th and SMI) and final serum C-peptide. Specifically, C-peptide significantly and positively correlated with Cs.Th (r= 0.63), Tb.Th (r= 0.45) and negatively with SMI (r= - 0.40). Furthermore, the table 3 shows a positive correlation between one of the IHC assessments (i.e. the PLIN1 score) and serum C-peptide (r= 0.52, Table 3). The Table also shows a negative correlation between neutrophils and C-peptide (r= −0.43). (S2 Fig.).

**Table 3:** Results of the nonparametric Kruskal-Wallis test for comparison among treatment groups of the radiologic parameters of microarchitecture and RANKL, as represented by the average and the standard deviation values, and the significance level (p) of the variation, also with correction test.

| Variables | CTR<sup>a</sup><br>(n=5) | D-CTR<sup>b</sup><br>(n=5) | C-PEP<sup>c</sup><br>(n=8) | p-value<br>(Kruskal-Wallis test) | p-value<br>(Mann-Whitney) | p-value<br>adjusted* | Post-hoc | CI 95% |  |
| --- | --- | --- | --- | --- | --- | --- | --- | --- | --- |
|  |  |  |  |  |  |  |  | Lower | Higher |
| Tb.Sp (mm) | 0.76±0.1 | 0.64±0.1 | 0.53±0.01 | 0.05 | a vs c, p=0.016 | 0.05 | a vs c, p=0.05 | -0.13 | 7.8 |
|  |  |  |  |  |  |  | b vs c, p=0.06 | -0.22 | 6.1 |
| CS.Th (mm) | 0.081±0.01 | 0.070±0.01 | 0.079±0.005 | 0.05 | a vs b, p=0.03 | 0.03 | / | / | / |
|  |  |  |  |  | b vs c, p=0.04 |  |  |  |  |
| Tb.Th (mm) | 0.122±0.001 | 0.111±0.001 | 0.125±0.007 | 0.02 | a vs b, p=0.03 | <0.001 | a vs c, p=0.04 | -6.1 | -0.01 |
|  |  |  |  |  | b vs c, p=0.01 |  |  |  |  |
| RANKL<br>(pg/mL) | 63±65 | 96±46 | 237±112 | 0.01 | a vs c, p=0.01 | 0.009 | b vs c, p=0.02 | -5.9 | -0.34 |
|  |  |  |  |  | b vs c, p=0.02 |  |  |  |  |

Note. Data are mean±SD. Tb.Sp Trabecular Separation; Cs.Th Crossectional thickness; Tb.Th Trabecular thickness; RANKL Receptor activator of nuclea factor kappa-B ligand. \*For baseline body weight.

S1 Table also reported the correlation between the final serum RANKL and several radiological and histological parameters of bone microarchitecture. RANKL positively correlated with the vBMD (r= 0.50). There was a negative correlation between Tb.Sp (assessed by both micro-CT and histomorphometry) and RANKL (r= - 0.61 and r= - 0.71, respectively)(S3 Fig.).

### 3.4 Micro-CT assessments

Three-dimensional images of femur epiphysis showed differences in trabecular microarchitecture among the various groups as represented in Fig.3. Analysis of the representative samples data indicate that C-peptide replacement therapy prevented the trabecular osteopenia.

**Figure 3:**
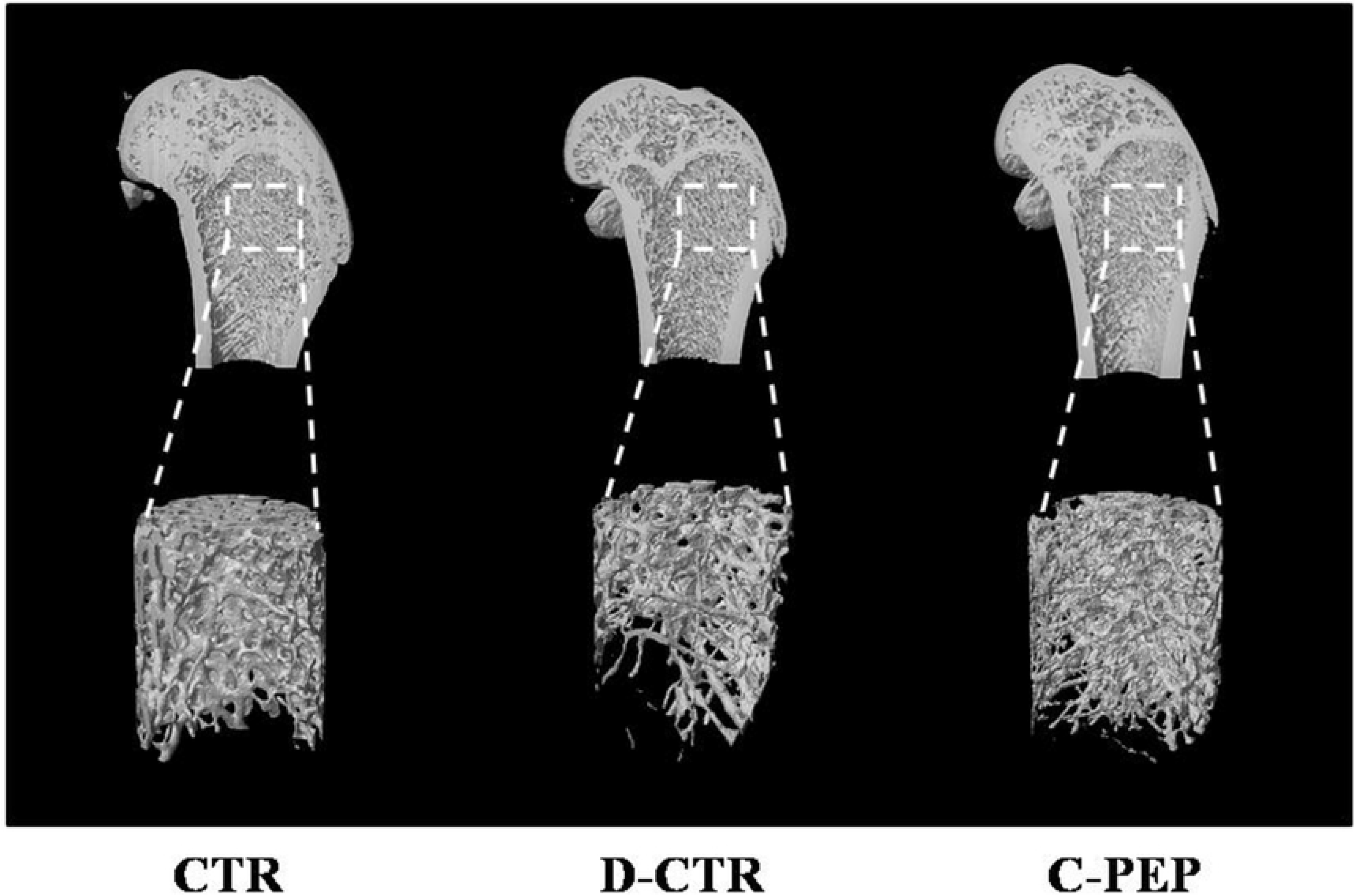
Three dimensional images of femur epiphysis among groups

Micro-CT evaluation of the femoral trabecular bone showed significant inter-group differences in Tb.Th (*p* = 0.022), also after adjustment (*p* <0.001, Table 3). The C-PEP group also showed lower Tb.sp than the other groups (adjusted *p* =0.05; CTR vs C-PEP *p* = 0.05 and D-CTR vs C-PEP p=0.06). There was a significant difference in Cs.Th, by Mann-Whitney test, between groups (CTR vs D-CTR *p* = 0.03 and D-CTR vs C-PEP p=0.04, Table 3), and this difference remained statistically significant after body weight adjustment (p=0.03, Table 3).

### 3.5 Histomorphometric parameters

After the treatment phase, neither of these parameters showed significant differences between groups. There was a significant correlation between the Tb.Th, assessed histologically, and several radiologic parameters (such as T.pm, B.pm, and Conn.D, see S1 Table).

### 3.6 Immunohistochemistry parameters

Both the CTR and C-PEP groups had more Runx-2 and PLIN1 positive cells (with a score of over 2) in comparison to the D-CTR group (p = 0.045 and p = 0.034, respectively, Fig.4).

**Figure 4:**
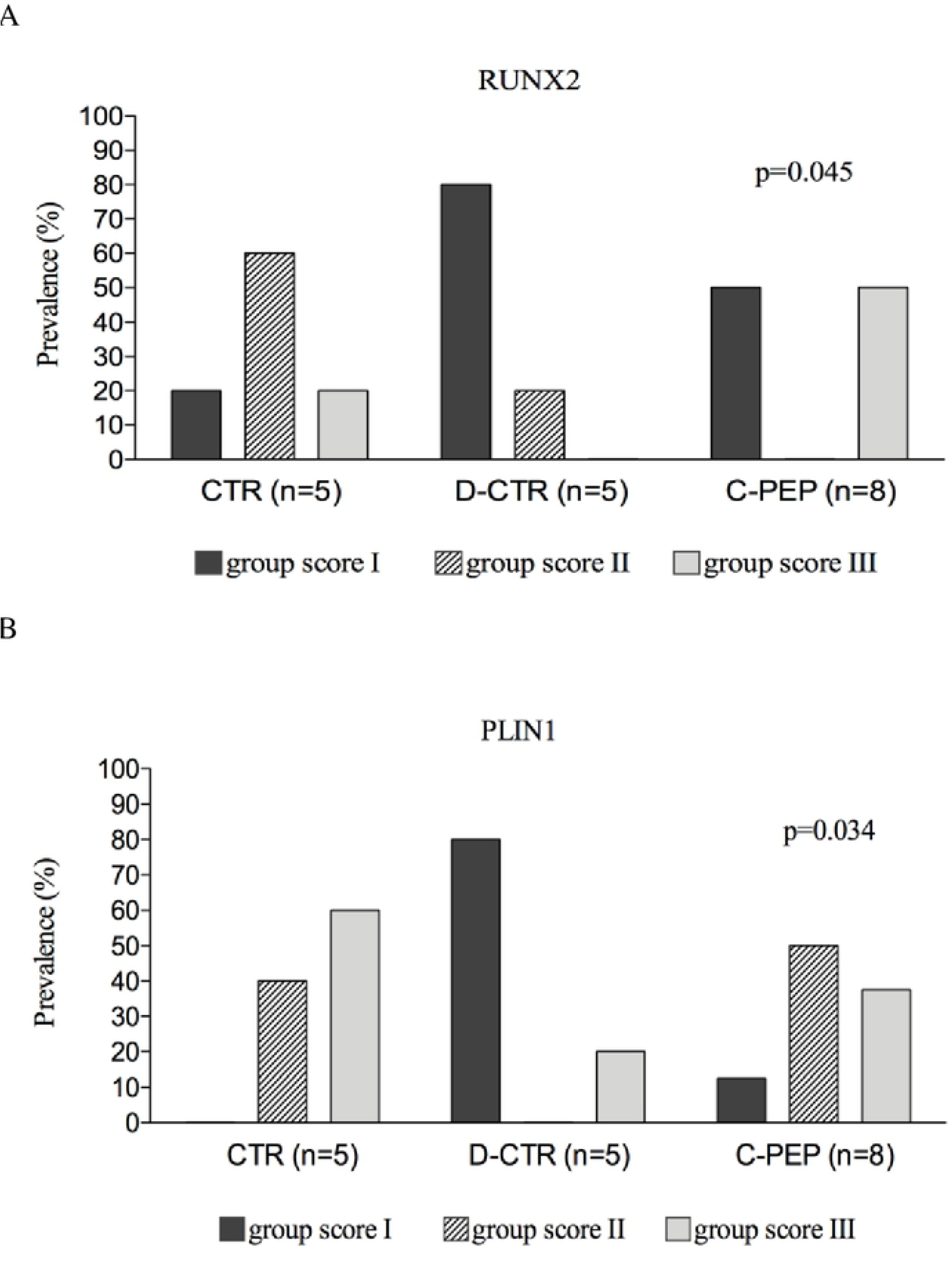
Hystological parameters score across intervention groups of (A) RUNX2 and (B) PLIN1

### 3.7 RANKL among groups

There was a significant difference in RANKL, by Mann-Whitney test, between groups (CTR vs C-PEP *p* = 0.01 and D-CTR vs C-PEP p=0.02, Table 3). After adjustment, C-PEP had a higher serum RANKL than D-CRT (D-CRT vs C-PEP, *p* =0.02, Table 3).

## 4. Discussion

Despite decades of research into the pathogenesis of diabetes, it is disappointing that the pathological pathways involved in the development of diabetic complications are still unclear. We hypothesized that a common pathophysiology for complications associated with T1DM exists, which also involves osteoporosis, that is a C-peptide deficiency. It has been demonstrated that T1DM complications develop even in individuals with good glycemic control [3,14]. Thus, C-peptide treatment could prevent bone microarchitecture damage and/or BMD deterioration induced by T1DM. The primary objective of this study was to test the effects of a 6-week regimen of continuous rat C-peptide infusion on bone of adult T1DM male rats, with C-peptide deficiency, by combining micro-CT imaging with histological testing.

These results, which are the first to demonstrate the effects of C-peptide replacement therapy *in vivo* on bone, indicate that treatment with the C-peptide could be beneficial in improving several bone microarchitecture parameters of the femur and, potentially, the bone mass of the tibia of T1DM rats. One initial interesting finding was that, at the end of the study, both the CTR and C-PEP groups exhibited a higher mean tibia weight than the D-CTR group (Table 2). We also found a significantly higher ash weigh from the left tibia in the C-PEP group, than in the CTR and D-CRT groups (Table 2).

In this study the quality of the trabecular bone was reflected by both the micro-CT and histological findings, which included a minimal set of variables to describe the effects of C-peptide on trabecular bone structure and, consequently, bone strength. Because patients with osteoporosis commonly sustain a fracture of the femur, in the present study we choose this anatomical region for all the trabecular assessments.

In our study, a significant difference in radiological and histological data was found between the groups. Compared to CTR rats, the femur of the D-CTR group had a 8 % (~ −0.01mm) lower Tb.Th and the C-PEP rats had a 2.5 % (~ +0.003 mm) higher Tb.Th (Table 3, p=0.022; adjusted p<0.001). The magnitude of the Tb.Th difference between the D-CTR and C-PEP rats (of ~ 0.015 mm) (Table 3), was the same as that observed in another study [39]. Animals in the C-PEP group also had a higher Cs.Th as well as a lower Tb.Sp than those in the D-CTR group (Table 3).

All these parameters are key measures in characterizing the three-dimensional structure of cancellous bone and correlate to bone strength [40]. Effective treatments may increase the trabecular bone by thickening the trabeculae. By its action on the osteoblasts [25], we hypothesise that C-peptide therapy led to the deposition of the new bone matrix in layers around the trabeculae, which thus became thick at the expense of the spaces between them. It is possible that the effects of C-peptide may also be exerted via blood glucose control. In fact, in line with previous reports of an increased whole body glucose utilization [19,22] and glucose transport in skeletal muscle [20], in our study, C-peptide treatment prevented the hyperglicemia seen in D-CTR (Fig.2). The co-administration of insulin and C-peptide in the conditions of their deficiency in diabetic rats led to the normalization of glucose homeostasis [41]. Further studies in humans are required to confirm this antidiabetic effect.

We found that both the CTR and C-PEP rats had, in femur sections, higher RUNX-2 proteins than the D-CTR group (Fig. 1A). This finding is in line with a study demonstrating that *Runx2* expression, which mediates the bone-forming action of Irisin, in turn triggers a global osteogenesis gene program [42]. Once expressed, this transcription factor drives osteoblasts to synthesize and secrete bone extracellular matrix, including type I collagen, bone sialoprotein and osteocalcin [42]. These actions could explain, at least in part, our finding of a high tibia weight in both the CTR and C-PEP groups.

All these findings were parallel with the changes observed, during the study, in the levels of serum C-peptide. At the end of the treatment period, serum C-peptide was higher in the C-PEP group than in the D-CTR group (Table 1, p=0.02) whereas it did not differ between the CTR and C-PEP groups (p = 0.09). The dose of C-peptide we used was probably insufficient to normalize the serum C-peptide. We, therefore, assume that, with a higher dose than that infused, we would have achieved this normalization.

Of note, the effect of C-peptide therapy on bone parameters sometimes exceeded those seen in the CTR group without C-peptide, which further highlights the potential efficacy of C-peptide as a pharmacological agent.

In line with previous studies (43,44), in the present investigation the D-CRT group exhibited similar microarchitectural parameter values to the CTR group (Tb.Sp ~ 0.70 mm in both CTR and D-CRT; Table 3). One study suggested that there was no obvious difference between the diabetic group and the control group at 4 weeks and 8 weeks in term of femur assessments, but a significant bone loss occurs in STZ-induced diabetes after 12 weeks [45]. This could also explain the lack of difference in vBMD between the rat groups as well as the bone histological parameters. Of course, histological methods only provide limited two-dimensional information, which is inadequate for the representation of comprehensive three-dimensional trabecular bone structures. A longer observation time would however have led to better observations of both the radiological and histological changes in the trabecular bone in T1DM rats.

In one study [28], which investigated the effects of amylin replacement therapy on bone loss in STZ-induced diabetic rats, the authors found that STZ-treated rats underwent a lower total BMD than nondiabetic control rats, after only 40 days of diabetes. Also considering the nominal duration of the osmopump (6 weeks), we designed this study choosing a 6 weeks study duration.

In our study, in histological samples from the femur of both C-PEP and CTR animals, we found a higher PLIN1 protein than in the D-CTR group. PLIN1 is known to be expressed only in adipocytes and macrophages [37]. PLIN1 plays a key role in the lipolysis of intracellular lipid deposits [37] and there may be significant associations between proteins that regulate fat metabolism and skeletal integrity. In fact, in bone, matrix vesicles are thought to arise by budding from the osteoblast plasma membrane containing high levels of cholesterol and other lipids, which ultimately result in the nucleation and propagation of mineral crystals [46]. However, Yamada et al. [46] studied a PLIN1 single-nucleotide polymorphism (SNP) and its relationship to BMD. They found a male-specific association, with the TT genotype having a greater BMD at several sites than the combined CC and CT genotypes. Future research is necessary to confirm the role of PLIN1 in macrophages differentiation as well as bone biology.

Another intriguing findings was that C-peptide administration prevented the red cells and haemoglobin reduction seen in D-CRT (Table 2). Thus, a potential effect of C-peptide on bone marrow could be assumed. There is a close relationship between haematopoiesis and bone formation. Anaemia may be more common in diabetes [48]. Haemoglobin levels are positively associated with BMD in the elderly, and anaemia is one of the risk factors for bone loss [49]. Mice deficient in Cbfa1/Runx2, a transcription factor crucial for osteoblast progression, did not develop osteoblasts and had empty bone marrow [50]. Osteoblasts are located at the endosteal bone surface just next to the hematopoietic stem cells (HSCs) in the bone marrow [51]. By producing several growth factors, osteoblasts increase the number of HSCs [52].

Through its effects on osteoblasts [25], we hypothesise that C-peptide, could, in turn, stimulate bone marrow-derived cells differentiation. Of course, we cannot confirm that this effect improves the trabecular bone. All these results need confirmatory research.

Another interesting finding relate to RANKL. In our study, C-PEP had a higher serum RANKL than D-CRT (Table 3). Since RANKL is expressed preferably by undifferentiated osteoblasts, this finding could reflect an increase in the number of active osteoblasts with the C-peptide therapy, as seen with Teriparatide [53]. In line with this concept, we found a negative correlation between Tb.Sp (assessed by both micro-CT and histomorphometry) and RANK-L (S1Table).

In our study C-peptide therapy was associated with both a high serum phosphorus and calcium concentration (Table 1). Rats received the same feeding schedule, thus, as observed with risedronate [54], our finding may confirm the direct role of C-peptide in the high bone turnover rate.

Although it is important to highlight the role of bone collagen in providing the matrix for bone mineralization and conferring bone elasticity, we found no association between COLIA1 or BSP and C-peptide concentration. Nor did we found a significant difference of COLIA1 or BSP, histologically assessed, between groups. Again, the study duration and/or the bone district analysed, likely affected these results.

In our study, other limitations should be acknowledged. We used a STZ-injected male rats model, however the effects in females could be different, thus the applicability of these results could be limited and could have affected our results.

Unfortunately, due to an electrical problem and the consequent partial loss of rats organs and tissues, we did not perform tibia measurements of the bone microarchitectural parameters. However, the potential clinical significance of C-peptide in osteoporosis is highlighted by all the present results.

## 5. Conclusions

This study, for the first time, demonstrated that the diabetic rats receiving rat C-peptide had higher quality of trabecular bone, represented by higher trabecular thickness, lower trabecular spaces and higher RUNX-2 protein in the femur, than diabetic rats not receiving this replacement therapy. In addition, C-peptide therapy increased the tibia weight. Finally, C-peptide administration prevented both the red cells and haemoglobin reduction seen in diabetic rats. Human studies are necessary to confirm the antidiabetic effect of C-peptide. However, in line with previous cross-sectional investigations carried out in postmenopausal women as well as *in vitro* observations, this study suggests the role of C-peptide in the prevention of bone damage, at least in T1DM.

## Funding

This research did not receive any specific grant from funding agencies in the public, commercial, or not-for-profit sectors.

## Declaration of interest

None.

## Acknowledgement

We are grateful to Dr John Wahren, from Karolinska Institutet, Stockholm, Sweden, for providing Rat C-Peptide.

## Credit Author Statement

Study design, data interpretation, drafting manuscript: T.M. and A.P.; Study conduct and data collection: S.M., C.R., V.M., S.N., M.G., F.B., M.S., M.R., L.D., M.T.V., M.D.; T.M., S.M., C.R., S.N., L.D.C., M.T.V., M.D. take responsibility for the integrity of the data analyses; revising manuscript content: V.M., D.B., V.M., L.D.C, S.R., S.G., C.G.; Approving final version and submitted version of manuscript: All authors

## Supporting Information

**S1 Table.** Bivariate correlations between serum C-peptide and several clinical, radiologic and histological parameters in rats

**S1 Fig.** Body weight change during the study

**S2 Fig.** Bivariate correlation between serum C-peptide and several radiologic and histologicalparameters in rats

**S3 Fig.** Correlation between RANKL and vBMD

